# Early hippocampal high-amplitude rhythmic spikes predict post-traumatic epilepsy in mice

**DOI:** 10.1101/2024.04.05.588288

**Authors:** Tyler Shannon, Noah Levine, Rina Dirickson, Yiyun Shen, Christopher Cotter, Yoon-Jae Yi, Noora Rajjoub, Fernando Pardo-Manuel de Villena, Olga Kokiko-Cochran, Bin Gu

**Affiliations:** Department of Neuroscience, Ohio State University, Columbus, USA; Electrical and Computer Engineering Program, Ohio State University, Columbus, USA; College of Veterinary Medicine, Ohio State University, Columbus, USA; Institute for Behavioral Medicine Research, Ohio State University, Columbus, USA; Department of Genetics, University of North Carolina, Chapel Hill, USA; Lineberger Comprehensive Cancer Center, University of North Carolina, Chapel Hill, USA; Chronic Brain Injury Program, Ohio State University, Columbus, USA

## Abstract

Oscillations, a highly conserved brain function across mammalian species, are pivotal in brain physiology and pathology. Traumatic brain injury (TBI) often leads to subacute and chronic brain oscillatory alterations associated with complications like post-traumatic epilepsy (PTE) in patients and animal models. Our recent work longitudinally recorded local field potential from the contralateral hippocampus of 12 strains of recombinant inbred Collaborative Cross (CC) mice alongside classical laboratory inbred C57BL/6J mice after lateral fluid percussion injury. In this study, we profiled the acute (<12 hr post-injury) and subacute (12-48 hr post-injury) hippocampal oscillatory responses to TBI and evaluated their predictive value for PTE. We found dynamic high-amplitude rhythmic spikes with elevated power density and reduced entropy that prevailed during the acute phase in CC031 mice who later developed PTE. This characteristic early brain oscillatory alteration is absent in CC031 sham controls or other CC and reference C57BL/6J strains that did not develop PTE after TBI. Our work provides quantitative measures linking early brain oscillation to PTE at a population level in mice under controlled experimental conditions. These findings will offer insights into circuit mechanisms and potential targets for neuromodulatory intervention.

## Introduction

Traumatic brain injury (TBI) causes various neurological sequela, such as sleep disruption, cognitive impairments, psychiatric complications, and recurrent seizures. Among them, post-traumatic epilepsy (PTE) emerges as one of the most prevalent and debilitating post-injury disorders. PTE significantly affects individuals’ quality of life and is associated with increased rate of unfavorable functional outcomes and mortality^1^. The higher risk of epilepsy and its related comorbidities in TBI underscores the critical need for novel resources and tools to identify individuals with TBI at risk for PTE. Despite extensive research into identifying biomarkers for TBI outcomes and epilepsy, there remains a paucity of studies investigating specific biomarkers for post-traumatic epileptogenesis.

Electroencephalography (EEG) is a widely used noninvasive neurodiagnostic tool for examining brain oscillatory function in both TBI and epilepsy. Abnormal EEG is common and heterogeneous following TBI, as early studies yield confilicing findings recarding its predictive utility for PTE^2^. Many descriptive and quantitative EEG alterations have been documented following TBI, including early post-traumatic seizures, high amplitude sharp waves, high-frequency oscillation (HFO), repetitive HFOs and spikes, sleep spindle duration, and EEG power change in both directions^2,3^. Retrospective case-controlled studies suggest some of these quantitative EEG features hold promise as predictive indicators for PTE^4-7^. However, these patients studies are face limitions such as poorly controlled environments, bias toward severe injury in participant selection, and insufficient consideration of patients’ genetics, demographic information, comorbid conditions, or medication history^8^. Though animal models offer greater control over environmental variability, early animal studies of TBI commonly use rodent models with fixed genetic backgrounds, failing to capture the genetic diversity seen in humans^9,10^. For example, the genotype-phenotype relationships observed in a single genetic background may not be generalizable to other backgrounds, limiting the translational revevance of such findings^11^.

We recently conducted a comprehensive characterization of TBI-related outcomes across 12 strains of the Collaborative Cross mice (CC), a next-generation multi-parental mouse genetic reference panel, in conjunction with the commonly used classical laboratory inbred C57BL/6J (B6J) mice^12^. CC provides a population of genetically diverse mice that allows us to systematically study the acute, subacute, and chronic brain oscillatory responses to TBI and how this information relates to PTE in a controlled environment. Among the 13 mice strains we studied, CC031 ermerged as the sole mouse strain displaying frequent spontaneous seizures after a moderate brain injury induced by lateral fluid percussion injury (FPI)^12^. Leverating this novel dataset, we quantitatively analyzed acute (< 12 hr) and subacute (12-48 hr) post-traumatic oscillatory features from the epileptic CC031 mice (CC031-TBI) by comparing them to non-epileptic CC and B6J strains that do not develop PTE after experiencing the same injury. We further analyzed new data from sham-operated CC031 and B6J mice as critical controls for injury-dependent phenotypes. This expanding dataset enables us to delineate temporal brain oscillatory features associated with PTE in consideration of genetic diversity. A deeper understanding of the dynamic changes in brain oscillation post TBI hold promise for identifying biomarkers and developing therapeutic intervention for PTE.

## Methods

Methods are reported as Supplementary Material.

## Results

### Dynamic high-amplitude rhythmic spikes (DHRS) prevail in the contralateral hippocampus during the acute phase after TBI in mice with PTE

Unlike most early preclinical EEG studies in TBI, which focused on analyzing the EEG recordings collected days, weeks, and months after injury, our study targeted the acute time window (<12 hours) immediately after TBI—a crucial yet understudied period wherein early brain oscillatory responses may serve as biomarkers of PTE. Through manual inspection of the initail 48 hours of LFP recordings covering both acute (< 12 hr) and subacute (12 – 48 hr) phases after TBI, we identified distinct electrographic abnormalities characterized by 0.3–0.5 Hz high voltage rhythmic spikes, namely DHRS, exclusively during the acute phase in all CC031-TBI mice. DHRS were absent in non-epileptic strains subjected to TBI or Sham-operated B6J and CC031 controls (Fig. 1A-1C and Supplemental Fig.1). Synchronized video recordings confirmed that DHRS occurrences coincided with periods of rest in CC031-TBI mice, ruling out the possibility that movement artifacts like grooming, eating, or drinking cause DHRS. Moreover, these DHRS were temporally restricted, occurring solely during the acute phase (average latency to onset = 386 ± 64 min and average duration = 178 ± 50 min), with the brain oscillations resuming normality thereafter until the emergence of post-traumatic seizures (Supplemental Fig.2). Notably, DHRS were also oberseved in one CC031 mouse who did not exhibit PTS during the three recording sessions after TBI, though longer continuous recordings are required to prove the presence of seizures in this particular mouse. In sum, our findings demonstrate the prescence of DHRS with characteristic power features exclusively in the mouse strain predisposed to PTE, highlighting a refined time window immediately following brain trauma for their detection.

**Figure 1.**
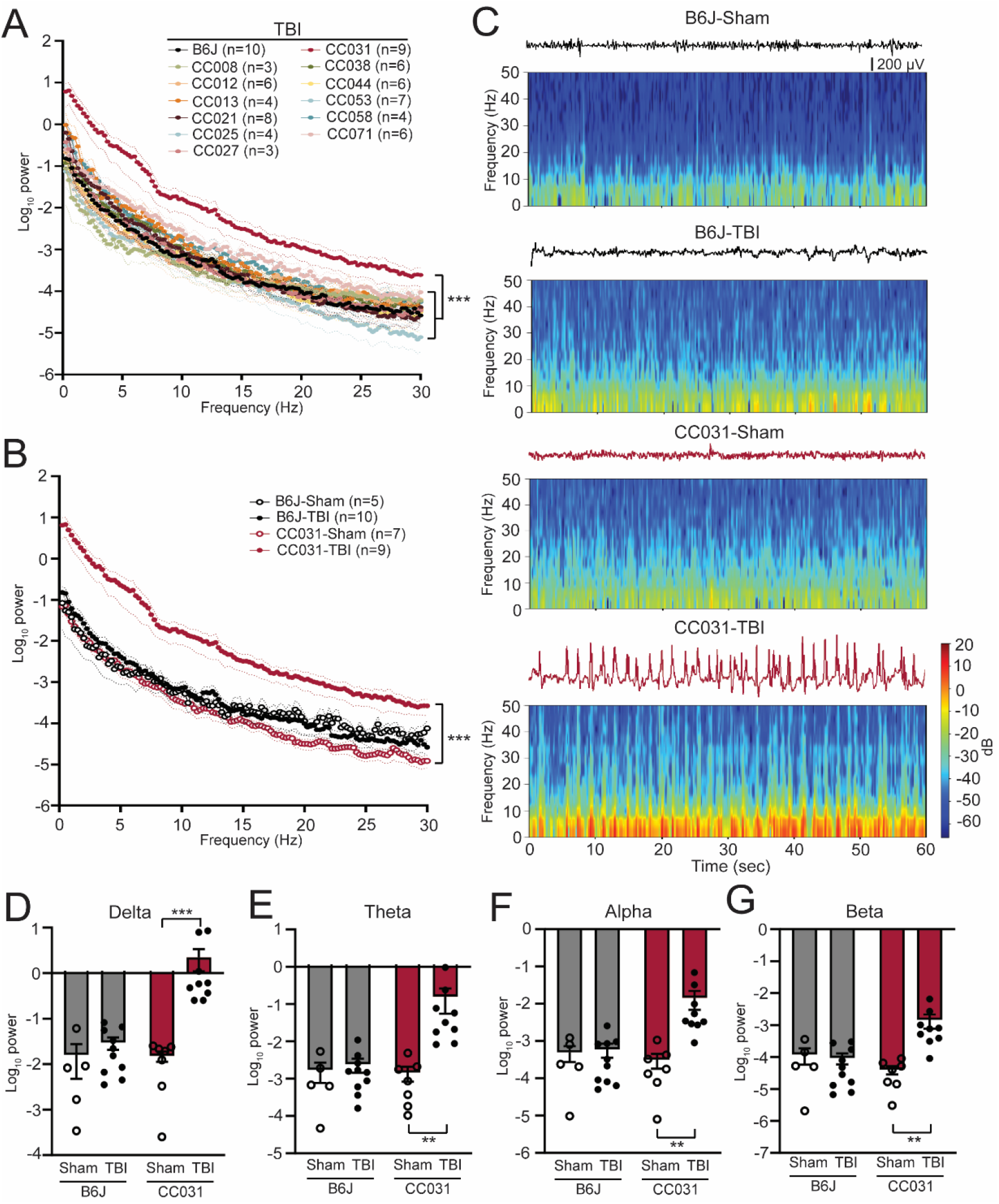
Acute contralateral hippocampal DHRS predict PTE. (A) Power spectral density analysis across 12 CC strains and B6J mice within 12 hr after TBI. Data are presented as mean ± SEM and analyzed using two-way ANOVA with post hoc Dunnett’s multiple comparison test, n = 3 – 10, ***P < 0.001 compared to B6J. (B) Power spectral density analysis of CC031 and B6J mice within 12 hr after TBI (replot) and their sham controls. Data are presented as mean ± SEM and analyzed using two-way ANOVA, ***P < 0.001, n = 5 – 10. (C) Representative EEG and spectrogram of CC031 and B6J mice within 12 hr after sham operation and TBI. (D-G) Accumulated power in (D) delta, (E) theta, (F) alpha, and (G) beta frequency bands in B6J and CC031 mice who were subject to sham operation or TBI. Data are presented as mean ± SEM and analyzed using two-way ANOVA with Šídák’s multiple comparisons test. n = 5 – 10, **P < 0.01 and ***P < 0.001.

### Quantitative brain oscillatory features predict PTE

We next quantitatively assessed the energy of DHRS by computing power spectral density using a randomly selected artifact-free two-minute time window during each phase, ensuring adequate capture of DHRS electrographic characteristics. During the acute phase, CC031-TBI mice exhibited a significant ubiquitous power elevation across the 0–30 Hz frequency domains compared to non-epileptic mouse strains (Fig. 1A). Given the comparable LFP features between B6J and non-PTE-developing CC strains and the impetus to use classical laboratory inbred mice, subsequent analyses focused on comparing epileptic CC031-TBI mice with non-epileptic control groups, including CC031-sham, B6J-sham, and B6J-TBI. We found the power density increased across the delta, theta, alpha, and beta bands in CC031-TBI mice compared to non-epileptic control groups (Fig. 1B-1G). These findings underscore the potential of quantitative brain oscillatory features, particularly acute delta power, in discriminating PTE and hold implications for predictive modeling and therapeutic interventions.

In addition to power analysis, we computated time-resolved PAC, a physiologically relevant activity observed in various brain areas that links to cognition and epilepsy^13,14^. We found that cross-frequency coupling of delta and gamma oscillations (fP: 0.5–4 Hz/fA: 30–200 Hz) are dominant in all four groups of CC031-Sham, CC031-TBI, B6J-sham, and B6J-TBI mice (Fig. 2A). We then focused on analyzing the average coupling strength of the dominant delta/low gamma (fP: 0.5–4 Hz/fA: 30–70 Hz) and delta/high gamma (fP: 0.5–4 Hz/fA: 80–200 Hz) and found their coupling strengths are indifferent regardless of injury or strain (Fig. 2B-C). Despite similar coupling strength, we found the preferred phase of delta/gamma coupling is predominantly located in the rising phase of delta wave in CC031-sham control mice, while this preference shifted to the falling phase in CC031-TBI mice who later developed PTE (Supplemental Fig. 3). Besides linear measurement, we assessed the non-linear features like the randomness of LFP by computing approximate entropy (ApEn), sample entropy (SampEn), and spectral entropy (SpecEn). Our analysis revealed significantly lower randomness of LFP power and spectrum in CC031-TBI mice who developed PTE compared to non-epileptic CC031-Sham, B6J-Sham, and B6J-TBI controls (Fig. 2D-F and Supplemental Fig. 4). Collectively, these immediate brain oscillatory alterations associated with PTE offer novel biological insights toward how acute electrophysiological response determines the risk of PTE and suggest potential targets for PTE prevention through neuromodulation.

**Figure 2.**
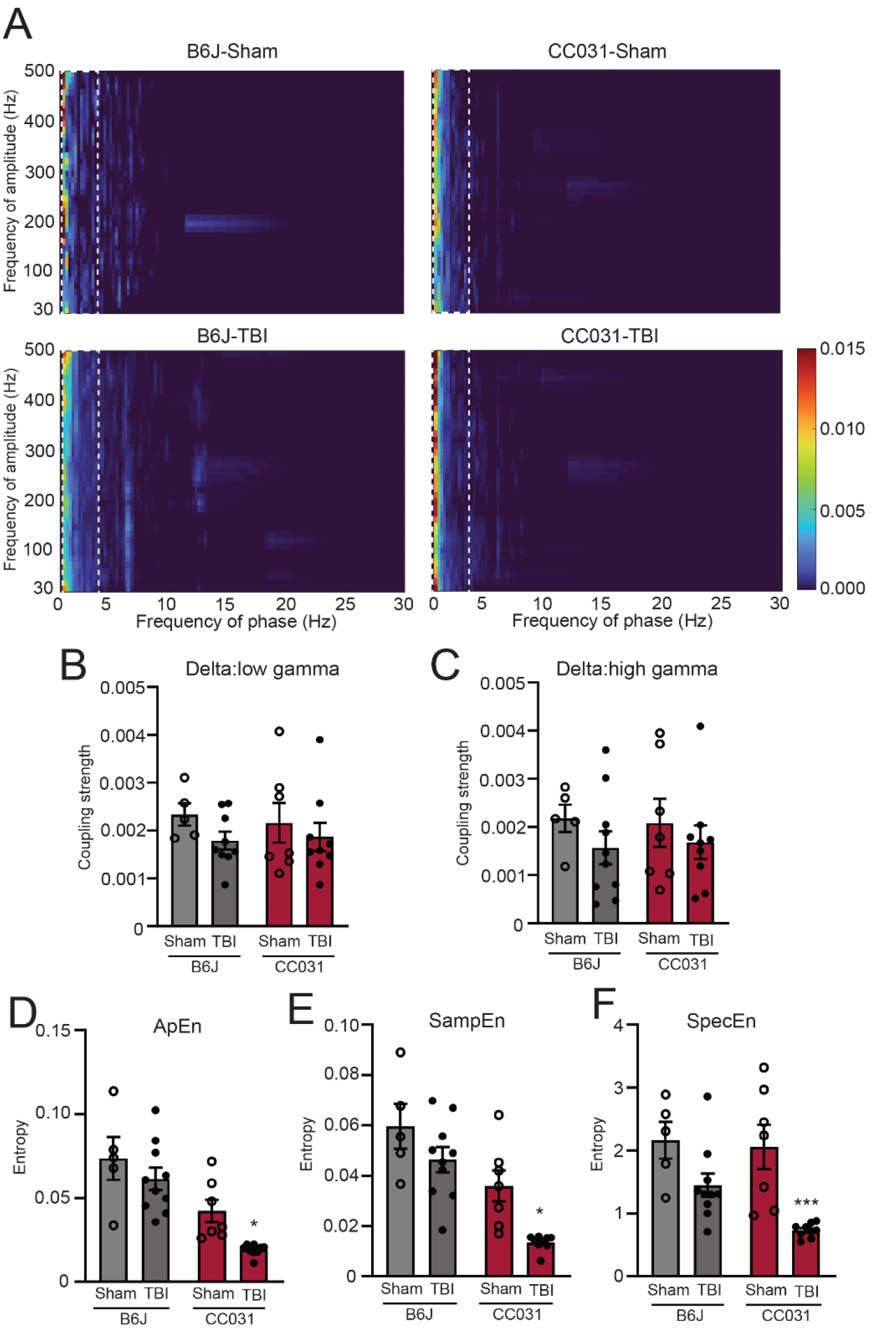
Lower early EEG entropy is dominant in the hippocampus of CC031 mice who develop PTE after TBI. (A) Average comodulogram of hippocampal PAC (fP: 0.5–30 Hz/fA: 30–500 Hz) in B6J-Sham, B6J-TBI, CC031-Sham, and CC031-TBI mice. The dashed rectangle denotes the dominant delta/gamma PAC (fP: 0.5–4 Hz/fA: 30–500 Hz). (B) and (C) Average coupling strength of delta/low gamma (fP: 0.5–4 Hz/fA: 30–70 Hz) and delta/high gamma (fP: 0.5–4 Hz/fA: 80–200 Hz, top row) PAC ranges. Analyses of (D) Approximate entropy (ApEn), (E) sample entropy (SampEn), and (F) spectral entropy (SpecEn) from the same time window. Data are presented as mean ± SEM and analyzed using two-way ANOVA with Šídák’s multiple comparisons test, *P < 0.05 and ***P < 0.001, n = 5 – 10.

## Discussion

Utilizing a novel mouse model of PTE within the CC panel, we conducted the first systematic EEG biomarker study after TBI in a genetically diverse mouse population under controlled experimental conditions. Our findings reveal that DHRS are present during an acute time window exclusively in mice who developed PTE after TBI. The absence of such abnormalities in other CC strains not exhibiting PTE after TBI suggests these oscillatory alterations are associated with immediate pathological responses of the brain to trauma that drives epileptogenesis. Clinically, our results hold significant implications on two fronts. Firstly, EEG forms a crucial component of the multimodal recordings of the intensive care unit (ICU), where patients are admitted after sustaining a TBI^15^. Both acute and subacute EEG have been increasingly collected as a routine procedure after TBI to facilitate diagnosis and indicate the prognosis of TBI^16^. Our findings align with a recent case-control study showing quantitative EEG features, including higher spectral power in the delta wave during early admission (within 3–5 days), can discriminate the risk of PTE after severe TBI^5^. Secondly, EEG provides a noninvasive real-time monitoring of the brain state and offers non-linear measurement of the dynamic changes of oscillations, reflecting the biological functions of the brain. These properties position EEG as a promising biomarker for PTE. Identifying early DHRS that precedes PTE also sheds light on the cellular mechanism of epileptogenesis after TBI. We speculate that DHRS may stem from a train of paroxysmal depolarization shifts in a large population of hippocampal CA1 pyramidal cells, which can be evoked by GABAa receptor inhibition, excessive glutamate release, or perturbed calcium homeostasis^17^.

Though DHRS offers a reliable biomarker in predicting PTE after FPI in CC mice, it is imperative to validate the generalizability of our findings by studying acute brain oscillation in other animal models of PTE. While the injuries were induced during the light cycle and most DHRS were observed during the subsequent dark cycle, the light/dark and circadian effects on these brain oscillatory biomarkers can be further investigated under reversed light/dark or constant light cycles. Further investigation using neuromodulatory approaches like phase-locked stimulation to suppress DHRS is required to ascertain a direct link between DHRS and PTE^18^. Inclusion of female mice is a critical next step in studying the sex effect on the brain oscillatory biomarkers of PTE. Despite these limitations, our study unveils early EEG characteristics preceding PTE, offering potential for improved risk stratification of patients, the development of new preventative and therapeutic modalities, and the prospect of ‘individualizing’ patient management based on their EEG findings.

## Authorship contributions

T Shannon: project management, performing experiments and data analysis; N Levine: code development, data curation, and analysis; R Dirickson: data curation and analysis; Y Shen: code development and data analysis; C Cotter: performing experiments; YJ Yi: performing experiments; N Rajjoub: performing experiments; F Pardo-Manuel de Villena: conceptualization and project management; O Kokiko-Cochran: conceptualization and writing; and B Gu: conceptualization, project management, conducting experiments, data analysis, and writing.

## Transparency, Rigor, and Reproducibility Summary

All CC strains were acquired from the Systems Genetics Core Facility at the University of North Carolina (UNC) at Chapel Hill between 2021 and 2023. B6J mice (#000664) were purchased from The Jackson Laboratory. Adult (3–6 months) male mice were used in this study. For TBI group, we used CC008 (n = 3), CC012 (n = 6), CC013 (n = 4), CC021 (n = 8), CC025 (n = 4), CC027 (n = 3), CC031 (n = 9), CC038 (n = 6), CC044 (n = 6), CC053 (n = 7), CC058 (n = 4), CC071 (n = 6), and B6J (n = 10) mice across 11 cohorts. For sham operated control group, we included CC031 (n = 7) and B6J (n = 5) mice over 3 cohorts. CC031 and B6J mice were randomly assigned to TBI and sham groups.

All experiments and analyses were performed by investigators who were blind to mouse strains and experimental groups. The Shapiro-Wilk normality test was performed to justify parametric statistical tests assuming Gaussian distribution. If data failed the normality test, we applied the appropriate non-parametric statistical tests instead. All data were presented as mean ± SEM. Unless otherwise noted, comparisons were analyzed using paired Student’s T-test or two-way ANOVA with Šídák’s post hoc test. P < 0.05 was considered statistically significant. Data and customized scripts are available from the corresponding authors upon request.

## Reference

1. Englander, J., Bushnik, T., Wright, J. M., Jamison, L.,Duong, T. T. Mortality in late post-traumatic seizures. Journal of neurotrauma 2009; 26, 1471–1477.

2. Perucca, P., Smith, G., Santana-Gomez, C., Bragin, A.,Staba, R. Electrophysiological biomarkers of epileptogenicity after traumatic brain injury. Neurobiol Dis 2019; 123, 69–74.

3. Haneef, Z., Levin, H. S., Frost, J. D., Jr.,Mizrahi, E. M. Electroencephalography and quantitative electroencephalography in mild traumatic brain injury. Journal of neurotrauma 2013; 30, 653–656.

4. Chen, Y., Li, S., Ge, W. et al. Quantitative epileptiform burden and electroencephalography background features predict post-traumatic epilepsy. Journal of Neurology, Neurosurgery & Psychiatry 2023; 94, 245–249.

5. Pease, M., Elmer, J., Shahabadi, A. Z. et al. Predicting posttraumatic epilepsy using admission electroencephalography after severe traumatic brain injury. Epilepsia 2023.

6. Reid, A. Y., Bragin, A., Giza, C. C., Staba, R. J.,Engel, J., Jr. The progression of electrophysiologic abnormalities during epileptogenesis after experimental traumatic brain injury. Epilepsia 2016; 57, 1558–1567.

7. Bragin, A., Li, L., Almajano, J. et al. Pathologic electrographic changes after experimental traumatic brain injury. Epilepsia 2016; 57, 735–745.

8. Rubinos, C., Waters, B.,Hirsch, L. J. Predicting and Treating Post-traumatic Epilepsy. Current Treatment Options in Neurology 2022; 24, 365–381.

9. Hajiaghamemar, M., Seidi, M., Oeur, R. A.,Margulies, S. S. Toward development of clinically translatable diagnostic and prognostic metrics of traumatic brain injury using animal models: A review and a look forward. Experimental Neurology 2019; 318, 101–123.

10. Brady, R. D., Casillas-Espinosa, P. M., Agoston, D. V. et al. Modelling traumatic brain injury and posttraumatic epilepsy in rodents. Neurobiology of disease 2019; 123, 8–19.

11. Sittig, Laura J., Carbonetto, P., Engel, Kyle A., Krauss, Kathleen S., Barrios-Camacho, Camila M.,Palmer, Abraham A. Genetic Background Limits Generalizability of Genotype-Phenotype Relationships. Neuron 2016; 91, 1253–1259.

12. Shannon, T., Cotter, C., Fitzgerald, J. et al. Genetic diversity drives extreme responses to traumatic brain injury and post-traumatic epilepsy. Experimental Neurology 2024, 114677.

13. Antonakakis, M., Dimitriadis, S. I., Zervakis, M. et al. Altered cross-frequency coupling in resting-state MEG after mild traumatic brain injury. Int J Psychophysiol 2016; 102, 1–11.

14. Liu, Y., McAfee, S. S., Guley, N. M. et al. Abnormalities in Dynamic Brain Activity Caused by Mild Traumatic Brain Injury Are Partially Rescued by the Cannabinoid Type-2 Receptor Inverse Agonist SMM-189. eNeuro 2017; 4.

15. Claassen, J., Taccone, F. S., Horn, P., Holtkamp, M., Stocchetti, N.,Oddo, M. Recommendations on the use of EEG monitoring in critically ill patients: consensus statement from the neurointensive care section of the ESICM. Intensive Care Med 2013; 39, 1337–1351.

16. Tewarie, P. K. B., Beernink, T. M. J., Eertman-Meyer, C. J. et al. Early EEG monitoring predicts clinical outcome in patients with moderate to severe traumatic brain injury. NeuroImage: Clinical 2023; 37, 103350.

17. Hotka, M.,Kubista, H. The paroxysmal depolarization shift in epilepsy research. Int J Biochem Cell Biol 2019; 107, 77–81.

18. Berényi, A., Belluscio, M., Mao, D.,Buzsáki, G. Closed-Loop Control of Epilepsy by Transcranial Electrical Stimulation. Science 2012; 337, 735–737.

