## Supplemental materials for "Early hippocampal high-amplitude rhythmic spikes predict post-traumatic epilepsy in mice"

**Supplemental methods**

**Animals**

All CC strains were acquired from the Systems Genetics Core Facility at the University of North Carolina (UNC) at Chapel Hill. B6J mice (#000664) were purchased from The Jackson Laboratory. All procedures were performed under the Institutional Animal Care and Use Committee (IACUC) of Ohio State University.

**Surgery**

Adult (3–6 months) male mice were anesthetized using isoflurane. Following a midline incision, a 3 mm diameter craniectomy was performed over the right hemisphere (coordinates from bregma: −0.5 mm AP and −2.5 mm ML) with careful attention to leaving the dura matter intact. A modified portion of a Luer-Loc (3 mm inner diameter) was positioned over the exposed dura and then secured over the craniectomy by cyanoacrylate adhesive and dental acrylic. Mice were returned to the home cage to recover overnight. One day after craniectomy, lateral FPI was delivered in isoflurane-anesthetized mice. Post-traumatic righting reflex time was assessed by the investigator as a surrogate measurement of the severity of the injury before the mouse was re-anesthetized using isoflurane. After FPI or sham injury, the Luer-Loc and adhesive were removed. A stainless steel bipolar recording electrode (P1 tech) was stereotaxically placed into the left dorsal hippocampus (coordinates from bregma: 2.0 mm AP, 1.6 mm ML, and 1.0 mm below dura). A ground electrode was fastened to a stainless steel screw on the skull above the left olfactory bulb. The electrodes were fixed, and the opening was closed using dental acrylic.

**Lateral FPI**

One day after craniectomy, all mice were anesthetized using isoflurane. Diffuse TBI was induced by delivering a 10–20 millisecond pulse of saline (1.2–1.5 atmospheres) onto the intact dura through the injury hub, which is equivalent to moderate TBI, using an FPI apparatus (Custom Design and Fabrication, Richmond, VA) as previously described. A separate group of age-matched CC031 mice were subject to sham operation and recorded as controls. Animals in the sham group were connected to the injury device for the same period of time; however, no fluid pulse was delivered.

**Video-local field potential (LFP) monitoring**

We performed time-locked video-LFP recordings across three time frames following injury: week 1, week 5–6, and week 9–10 post-injury. This design allows us to study early brain oscillatory biomarkers that can shape the later onset of PTE. During recording, the mouse was single-housed under diurnal conditions of 12/12 light and dark cycles. The recording started ~3 hr after completion of TBI to allow the animal to recover and regain voluntary activity from injury and surgery. The 24/7 continuous recordings were performed using a 12-channel acquisition system (Data Science International). Ponemah 5.5 software was used for data acquisition and video synchronization (infrared-enabled for night vision). The mouse can freely roam within the housing cage via a tethered system connected to a commutator. LFP recordings were sampled at a rate of 2000 Hz.

**Signal processing and analysis**

All LFP data were filtered using an $8^{th}$-order notch filter of 60 Hz before analysis. The power spectrum and phase-amplitude coupling were analyzed using *Brainstorm* and customized MATLAB algorithms. The spectrogram, phase preference, and entropy were analyzed and visualized using MNE and Tensorpac packages in Python. A random two-minute time window was selected during specific time frames of 3–12 hr (acute) and 12–48 hr (subacute) after TBI. The length of the time window was selected to reflect the patterns and features of LFP while considering the computational cost. For the power spectrum, the LFP data were firstly transformed using Morlet Wavelet with 3 sec time resolution and 0.5 Hz step size. The data were further divided into particular wave bands for further quantification: delta (0.5–4 Hz), theta (4–8 Hz), alpha (8–12 Hz), and beta (12–30 Hz). The performance of the PTE prediction model was evaluated using the receiver operating characteristic (ROC) and the area under the curve (AUC) ROC analysis. The spectrogram scale was adjusted to reveal details of the sample with the lowest power. For phase-amplitude coupling (PAC), the time-resolved PAC was computed using a 4-second sliding time window. MATLAB generated the average comodulogram in the frequency ranges for phase (fP, 0.5–15 Hz) and frequency for amplitude (fA, 30–200 Hz). The coupling strength of delta/low gamma with fP (0.5–4 Hz)/fA (30–70 Hz), delta/high gamma fP with (0.5–4 Hz)/fA (80–200 Hz), theta/low gamma with fP (4–8 Hz)/fA (30–70 Hz), and theta/high gamma with fP (4–8 Hz)/fA (80–200 Hz) was calculated. For phase preference,  we computed target fP of 0.5–4 Hz, minimum fA of 30 Hz, and maximum fA of 100 Hz with a 4-second sliding time window using Tensorpac in Python. The average phase preference was computed and plotted in a polar histogram. Entropy was calculated using the app_entropy, samp_entropy, and spect_entropy functions of MNE in Python.

**Statistical analysis**

All experiments and analyses were performed by investigators who were blind to mouse strains and experimental groups. The Shapiro-Wilk normality test was performed to justify parametric statistical tests assuming Gaussian distribution. If data failed the normality test, we applied the appropriate non-parametric statistical tests instead. All data were presented as mean ± SEM. Unless otherwise noted, comparisons were analyzed using paired Student's T-test or two-way ANOVA with Šídák's *post hoc* test. $P < 0.05$ was considered statistically significant.

**Supplemental figures 1–4**

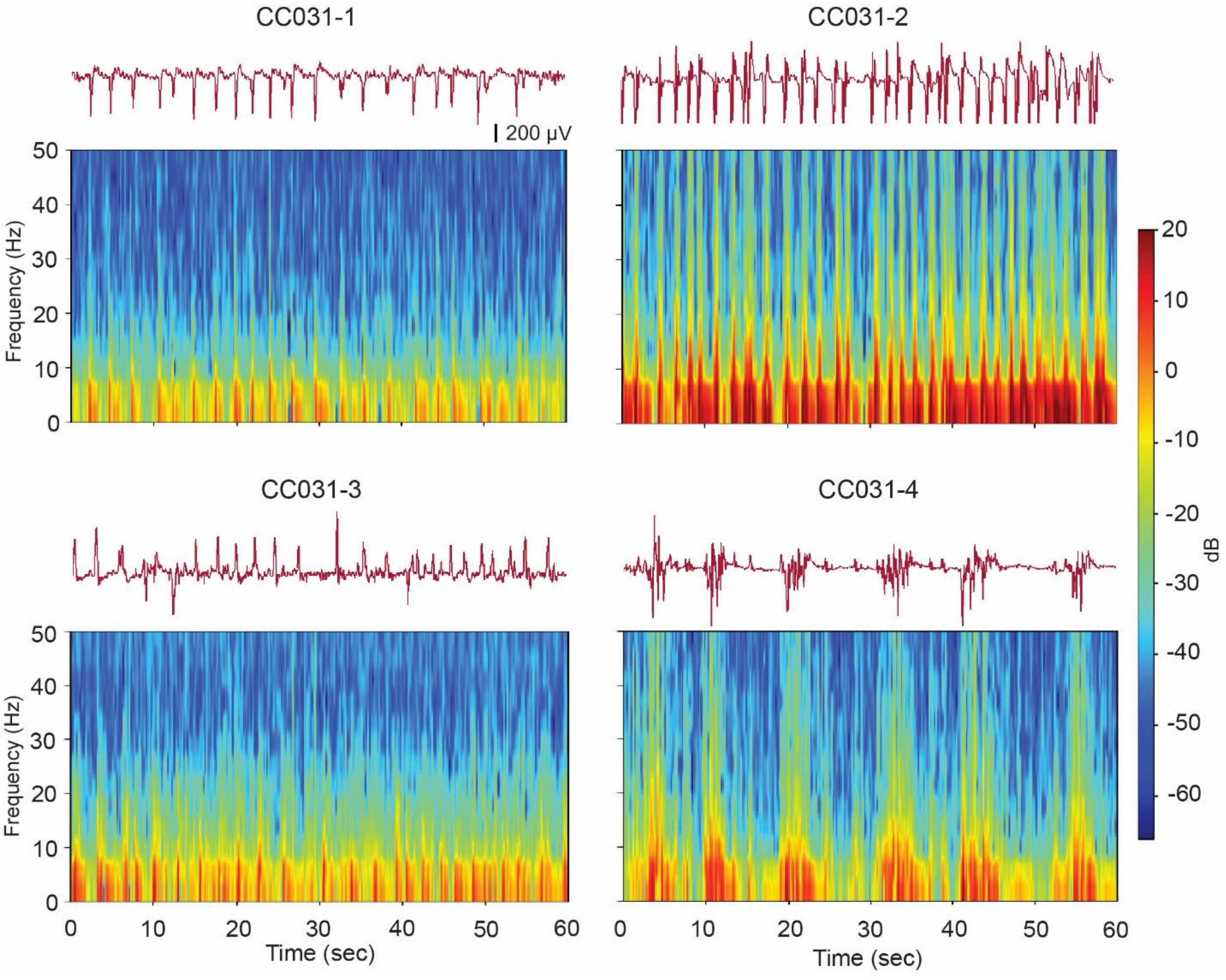


**Supplemental Figure 1. Representative hippocampal LFP traces and spectrograms of acute DHRS in CC031-TBI mice.** DHRS in CC031-1, CC031-2, and CC031-3 mice are characterized by 0.3–0.5 Hz high voltage rhythmic spikes. CC031-4 showed a unique DHRS pattern with bouts of 5 s spindles at a ~10 s interval.

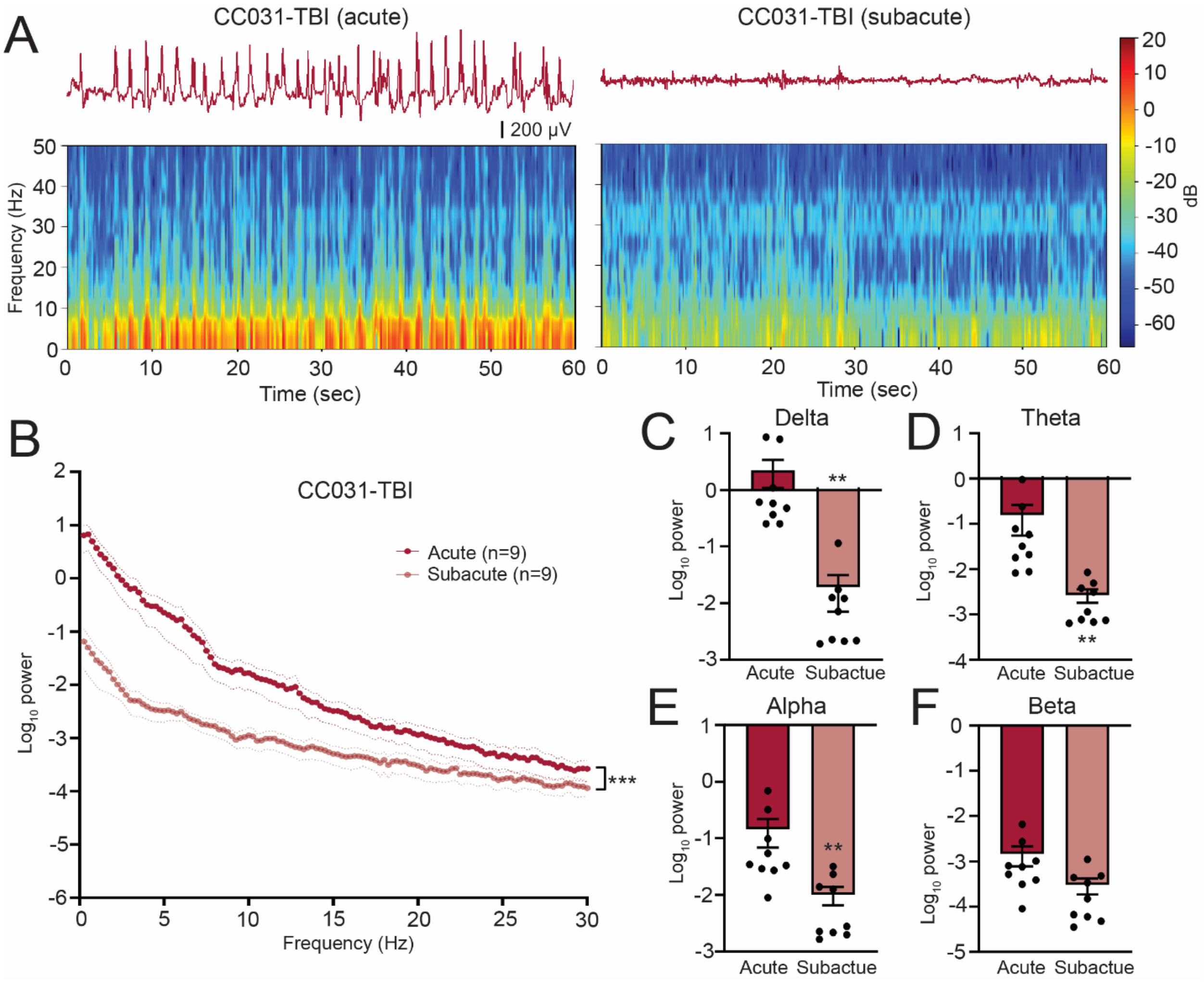


**Supplemental Figure 2. Hippocampal LFP resumes normality 12 hr after TBI in CC031 mice.** (A) Representative LFP traces and spectrograms during acute and subacute phases in a CC031-TBI mouse. (B) Power spectral density analysis of CC031-TBI mice during acute vs. subacute phases. Data are presented as mean ± SEM and analyzed using two-way ANOVA, ***$P < 0.001$. (C–F) Accumulated power in (C) delta, (D) theta, (E) alpha, and (F) beta frequency bands of CC031-TBI mice comparing acute vs. subacute phases. Data are presented as mean ± SEM and analyzed using Wilcoxon matched-pairs signed rank test, **$P < 0.01$.

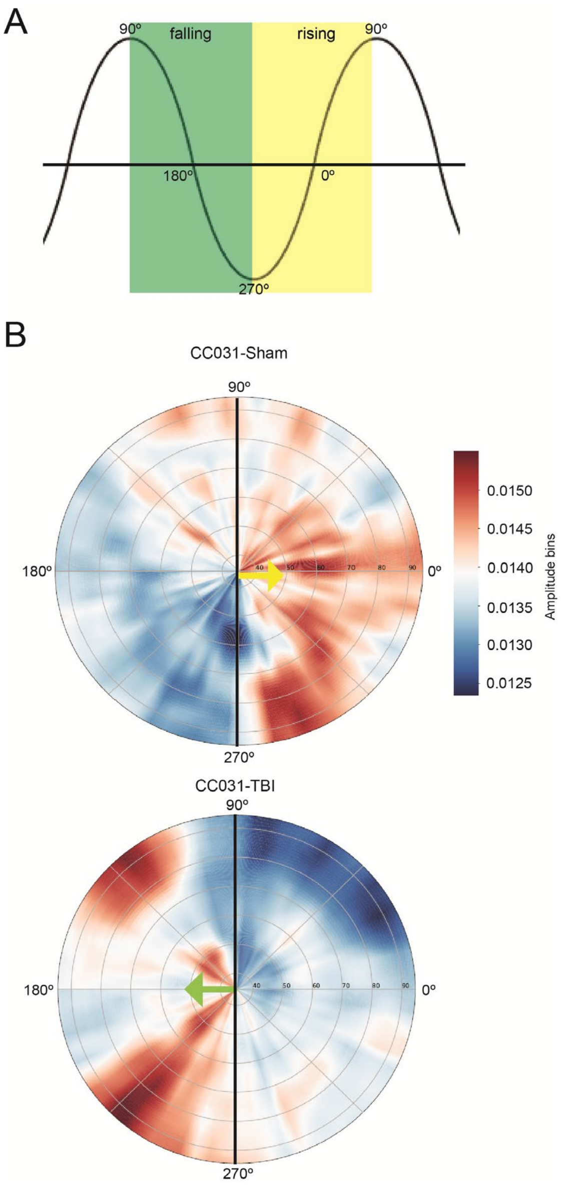


**Supplemental Figure 3. Phase preference of delta/gamma coupling is opposite in CC031-sham vs. CC031-TBI.** (A) Schematic of phase angle. The green square denotes the falling phase. The yellow square indicates the rising phase. (B) The angle represents the delta frequency (0.5–4 Hz) phase, and the radial axis represents different gamma frequencies (30–100 Hz) for the amplitude signal. The color depicts the average amplitude value of a given frequency inside the corresponding phase bin. Gamma oscillation (30–100 Hz) preferably riding on the rising phase of the delta wave in CC031-sham mice while on the falling phase of the delta wave in CC031-TBI mice.

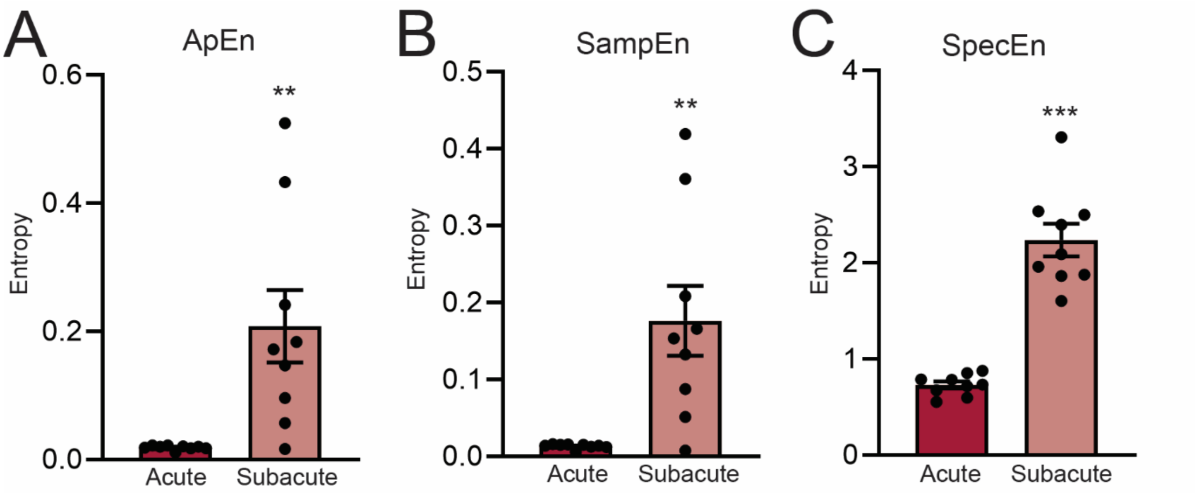


**Supplemental Figure 4. LFP entropy in the hippocampus of CC031 mice increased on 1 DPI compared to 0 DPI.** (A) Approximate entropy (ApEn), (B) sample entropy (SampEn), and (C) spectral entropy (SpecEn) of CC031-TBI mice were analyzed by comparing acute vs. subacute. Data are presented as mean ± SEM and analyzed using paired Student's t-test, **P < 0.01; ***P < 0.001.
